# Celloscope: a probabilistic model for marker-gene-driven cell type deconvolution in spatial transcriptomics data

**DOI:** 10.1101/2022.05.24.493193

**Authors:** Agnieszka Geras, Shadi Darvish Shafighi, Kacper Domżał, Igor Filipiuk, Łukasz Rączkowski, Hosein Toosi, Leszek Kaczmarek, Łukasz Koperski, Jens Lagergren, Dominika Nowis, Ewa Szczurek

## Abstract

Spatial transcriptomics maps gene expression across tissues, posing the challenge of determining the spatial arrangement of different cell types. However, spatial transcriptomics spots contain multiple cells. Therefore, the observed signal comes from mixtures of cells of different types. Here, we propose an innovative probabilistic model, Celloscope, that utilizes established prior knowledge on marker genes for cell type deconvolution from spatial transcriptomics data. Celloscope outperformed other methods on simulated data, successfully indicated known brain structures and spatially distinguished between inhibitory and excitatory neuron types based in mouse brain tissue, and dissected large heterogeneity of immune infiltrate composition in prostate gland tissue.

## Background

Gene expression is crucial for characterizing tissue functionality in both normal and abnormal conditions. For this reason, various high-throughput sequencing technologies measuring gene expression were developed. In contrast to standard bulk and single cell RNA sequencing (scRNA-seq) technologies (1–4), the groundbreaking spatial transcriptomics (ST) technology (5) enables to investigate the tissue functionality in an unprecedented, ultra-high, spatial resolution. ST captures RNA-sequencing measurements in multiple distinct spots localized on the examined tissue, yielding genes versus spots expression matrices.

Each ST spot contains an unknown number of cells, typically ranging between 10 and 100 cells (5, 6). Therefore, the observed signal inevitably conveys information about mixtures of cells of various types. Consequently, a number of computational methods for deconvolution of cell types in ST spots were proposed. These methods require additional scRNA-seq measurements, ideally from the same tissue (7–14). To perform the deconvolution, in the first step, the additional scRNA-seq data are used to compute gene expression profiles for the analyzed cell types. In the second step, the computed expression profiles are utilized to resolve the composition of cell types present at each ST spot. To his end, diverse statistical methods are applied, such as maximum a posteriori estimation in the case of Stereoscope (8) and maximum likelihood estimation in the case of RCDT (7). Cell2location (11), as a Bayesian hierarchical framework, allows for introducing factorized priors. However, the choice of hyperparameters may greatly influence its performance (11). An alternative statistical strategy for deconvolution is to use lognormal (10) or weighted linear regression (13), utilizing the pre-computed gene expression levels for cell types as explanatory variables. Finally, Tangram (14) learns a mapping function that transfers scRNA-seq (or single nuclei RNA-seq) expression to the tissue space measured using ST, so that spatial correlation across the genes between the two techniques is maximised. Integrating data obtained via two different experimental techniques such as scRNA-seq and ST may be prone to bias arising from cross-platform and batch effects, such as differences in the sequencing depth or only a partial overlap between sequenced cell types and genes. Moreover, such a reference data set measuring gene expression in cells of different types, ideally taken from exactly the same tissue sample, may not be accessible. This calls for methods designed for ST data that do not require an external reference in the form of scRNA-seq data.

To address these needs, here we present Celloscope – a novel method for decomposing cell type mixtures in ST spots, enabling to spatially map cell types in tissues examined with ST technology. Importantly, the innovative character of Celloscope involves using prior *qualitative* information on marker genes. Such marker genes are expected to be more highly expressed in their respective cell types compared to other types (15). There is a large body of expert knowledge on marker genes for different cell types, established by multiple independent studies (as of 09.04.2022, PubMed search for “cell type marker” phrase returned 67,327 entries). Celloscope can leverage this wealth of knowledge, and, as a consequence, is fully independent of scRNA-seq reference datasets. The expression level for each marker gene in each cell type is estimated as part of the model’s inference from ST data. Moreover, we estimate the admixture of additional, *a priori* unknown cell types.

The results of extensive experiments on simulated data demonstrated the excellent Celloscope’s performance. On top of that, Celloscope surpassed other methods that use scRNA-seq data as a reference to compute gene expression profiles in a simulation study. To further validate the correctness of the Celloscope’s assumptions and prove its usefulness, we applied it on data on the anterior section of the mouse brain. We also showed that Celloscope can be used to elucidate the cause of inflammation in a human prostate tissue (16). These results indicate that Celloscope successfully leverages biological knowledge of cell type marker genes to accurately decompose cell type mixtures in ST data.

## Results

### Celloscope model overview

We propose Celloscope, a novel Bayesian probabilistic graphical model of gene expression in ST data, which deconvolutes cell type composition in ST spots, and a method to infer model parameters based on a MCMC algorithm (Methods). Apart from gene expression measurements per each spot localized in the analyzed tissue, the ST data comes also with corresponding images of hematoxylin and eosin (H&E) staining of the same sample. The first step of the Celloscope’s pipeline is an analysis of those H&E images (Fig. 1A). In this step, the number of cells for each spot is estimated. Optionally, regions of interest in the tissue (e.g., an inflammation) are annotated by a pathologist, so that further analysis steps are restricted to those regions.

**Fig. 1.**
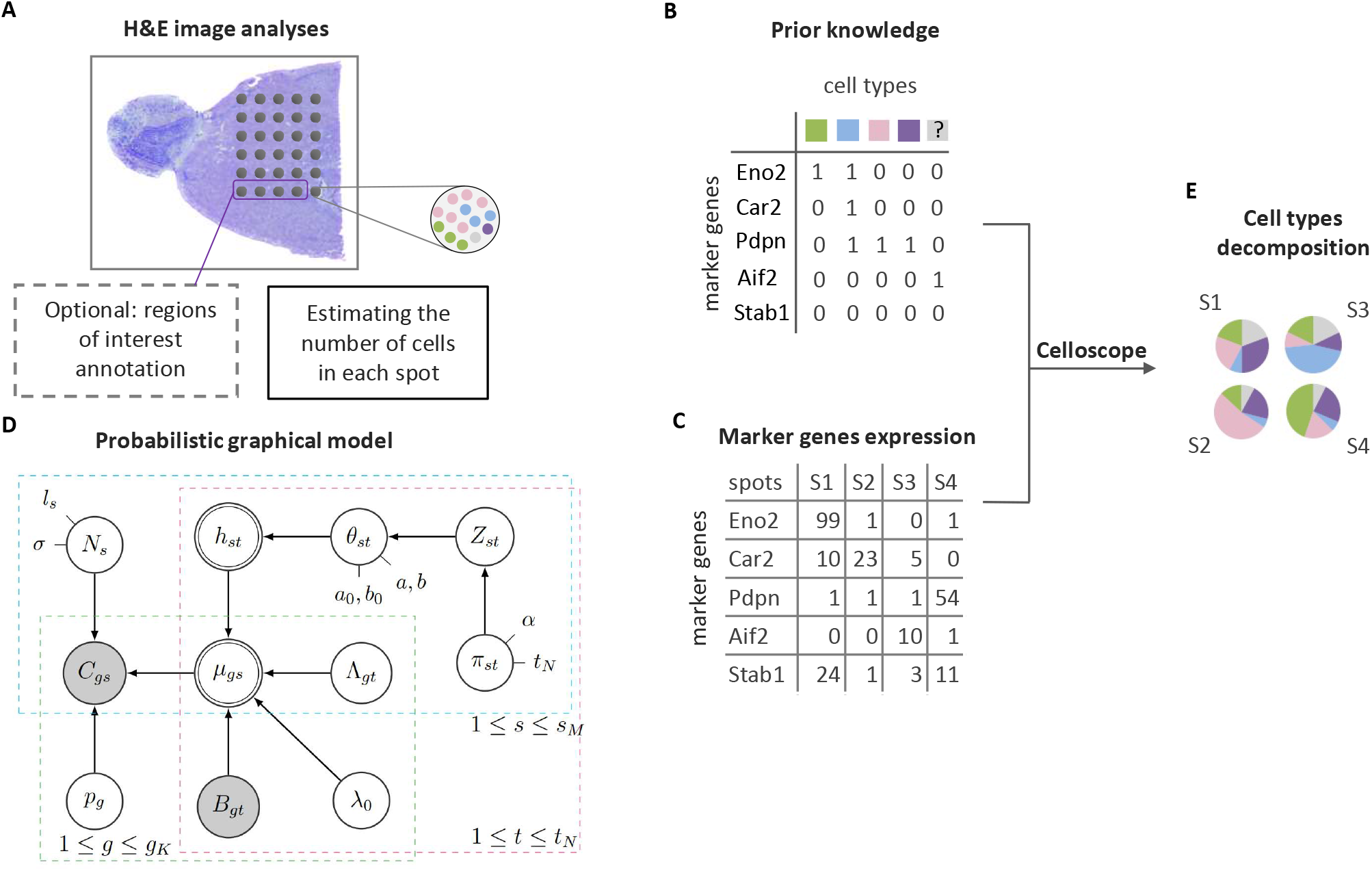
Celloscope overview. **A** The total number of cells for each spot is estimated based on a H&E image. Optionally, regions of interest are annotated. H&E image source: 10X Genomics. **B** Prior knowledge on marker genes is given as the model’s input, in a form of a binary matrix, together with ST data on marker gene expression in spots (**C**). **D** The graphical representation of Celloscope. Gray nodes correspond to the observed variables, double circled to deterministic ones, while the remaining nodes correspond to hidden variables. Arrows represent probabilistic dependencies. Model’s variables are described in Table 1 and hyperparameters in Table 2. **E** Cell type decomposition in each spot using Celloscope is performed via MCMC inference.

Prior knowledge on marker genes for considered cell types is encoded as a binary matrix (Fig. 1B). The types correspond to such cells that are expected to be found in the examined tissue or (optionally) in the selected area of interest. Additionally, we assume the presence of a *dummy type* accounting for novel or unknown cell types. The dummy type is characterised by zero marker genes.

The estimated cell counts and the binary matrix encoding prior knowledge on marker genes, together with the measured expression of the marker genes in each (selected) spot (Fig. 1C) constitute the input to the probabilistic model (Fig. 1D). The model assumes that the measured expression depends on the hidden cell type mixture in each spot. As the output, Celloscope returns proportions of cell types for each spot of interest (Fig. 1E).

### Celloscope’s results on simulated data prove exceptional performance of marker gene-based cell type deconvolution

To demonstrate excellent performance of Celloscope in marker gene-based cell type deconvolution, we tested the model on different setups of simulated data, for which we knew the ground truth about the underlying cell type composition, and therefore we were able to confront the true cell type proportions across spots with the model’s estimates. We consider four distinct simulation scenarios which differ by two key simulation parameters. The first simulation parameter is the average number of cell types present within each spot, with the *dense* scenario implying an increased number, and the *sparse* scenario denoting a deceased number of cell types per spot. The second simulation parameter accounts for the incorporation of the true values of the number of cells in each spot, which are either treated as *known* and given as observed variables to the model, or *as priors*, i.e., they are introduced as informative priors for the hidden variables that correspond to the number of all cells in each spot. For each scenario, 15 datasets were simulated, assuming 800 spots, 8 cell types and 149 marker genes. We computed the average absolute error across spots for each replicate.

Celloscope achieved excellent performance, with error levels between 0.01 and 0.03 (Fig. 2A), proving that knowledge of marker genes for cell types is sufficient for accurate cell type deconvolution in ST spots. The additional information on the number of cells in each spot increased the accuracy of the model’s prediction.

**Fig. 2.**
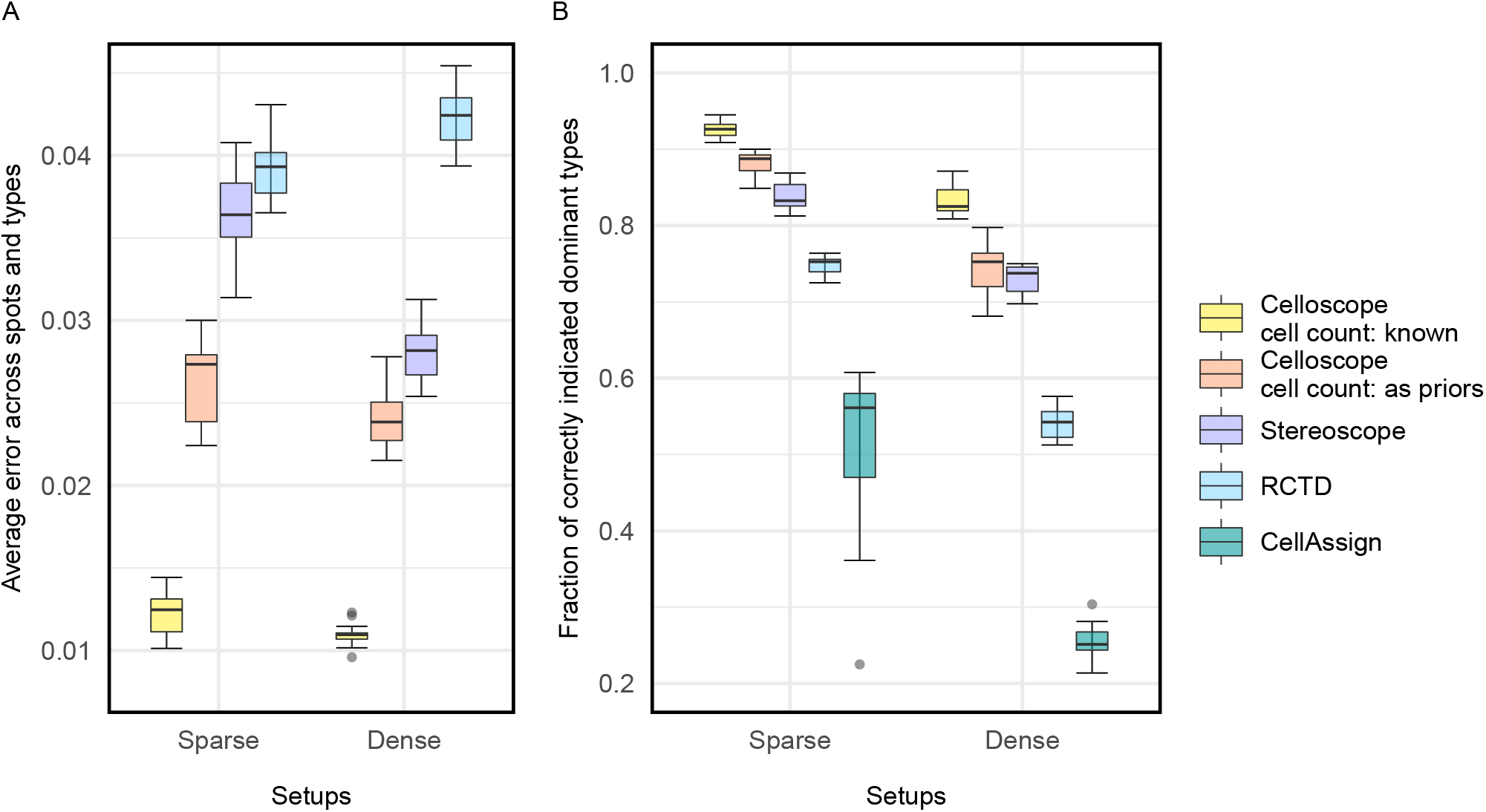
Excellent performance of Celloscope on simulated data. **A** Box-plots represent distributions of average absolute error (y-axis; computed using Equation 16) for different methods (colors) and data simulation scenarios (x-axis). Celloscope outperforms competing methods, Stereoscope (8) and RCTD (7) which rely on additional input regarding gene expression levels in different cell types. **B** Distribution of the fraction of correctly identified dominant cell types across spots (y-axis), for different methods (colors) and simulation scenarios (x-axis). Celloscope again shows a large advantage over other methods. Here, we compare also to CellAssign (17), indicating the benefit from decomposing a mixture of different cell types in ST spots.

#### Comparison to preceding approaches

We compared Celloscope’s performance with previously published Stereoscope (8) and RCTD (7). Both of these methods, unlike Celloscope, require a reference scRNA-seq dataset to compute gene expression profiles for the analysed cell types. Importantly, all considered methods were applied on exactly the same simulated datasets, for which the ground truth was known. Thus, we were in possession of the true marker gene expression levels across cell types, which were provided to RCTD and Stereoscope and used by these methods for the estimation of cell type proportions in simulated spots (see Additional File 1: Section S1 and S2 for the implementation details and run settings used for RCTD and Stereoscope, respectively). These values were not provided to Celloscope, but inferred. Therefore, RCTD and Stereoscope were given a head start as compared to our model.

Despite being given such an advantage, both RCTD and Stereoscope performed poorly compared to Celloscope (Fig. 2A). The observed much higher error for RCTD might occur due to the fact that this model uses Poisson distribution to model gene expression, in contrast to Celloscope and Stereoscope, that utilise the Negative Binomial distribution.

To evaluate the importance of the assumption of the presence of a mixture of cells of different types in ST spots and performing cell type decomposition by inferring propositions of cell types, in contrast to indicating only the dominant type, we compared our results to CellAssign (17) (run settings provided in Additional File 1: Section S3). Similarly to Celloscope, CellAssign uses a binary cell type marker matrix as model input. However, Celloscope was originally developed to assign cell types in scRNA-seq data, and as such it assumes that each observation refers to only one cell (of a given type). Therefore, we expect that CellAssign, applied to simulated ST data, will treat each spot as homogeneous and indicate the dominant type. For each spot, we checked if the true dominant cell type (the cell type characterised by the highest proportion) was in agreement with the inferred dominant type. Methods dedicated to ST and performing the cell type deconvolution performed the task of finding the dominant type significantly better than CellAssign (Fig. 2B), which confirms the key importance of accounting of the mixture of cell types in each ST spot.

### Celloscope localises cell types in agreement with known mouse brain structures

Celloscope was applied on mouse brain data (18) and was able to successfully indicate brain structures (Fig. 3A). Specifically, an analysis of spatial transcriptomics data on sagittal mouse brain slices (an anterior section) was performed. Convergence diagnostics using the Gelman-Rubin method showed high evidence for the convergence of our parameter sampling procedure (Additional File 1: Figure S1 A).

**Fig. 3.**
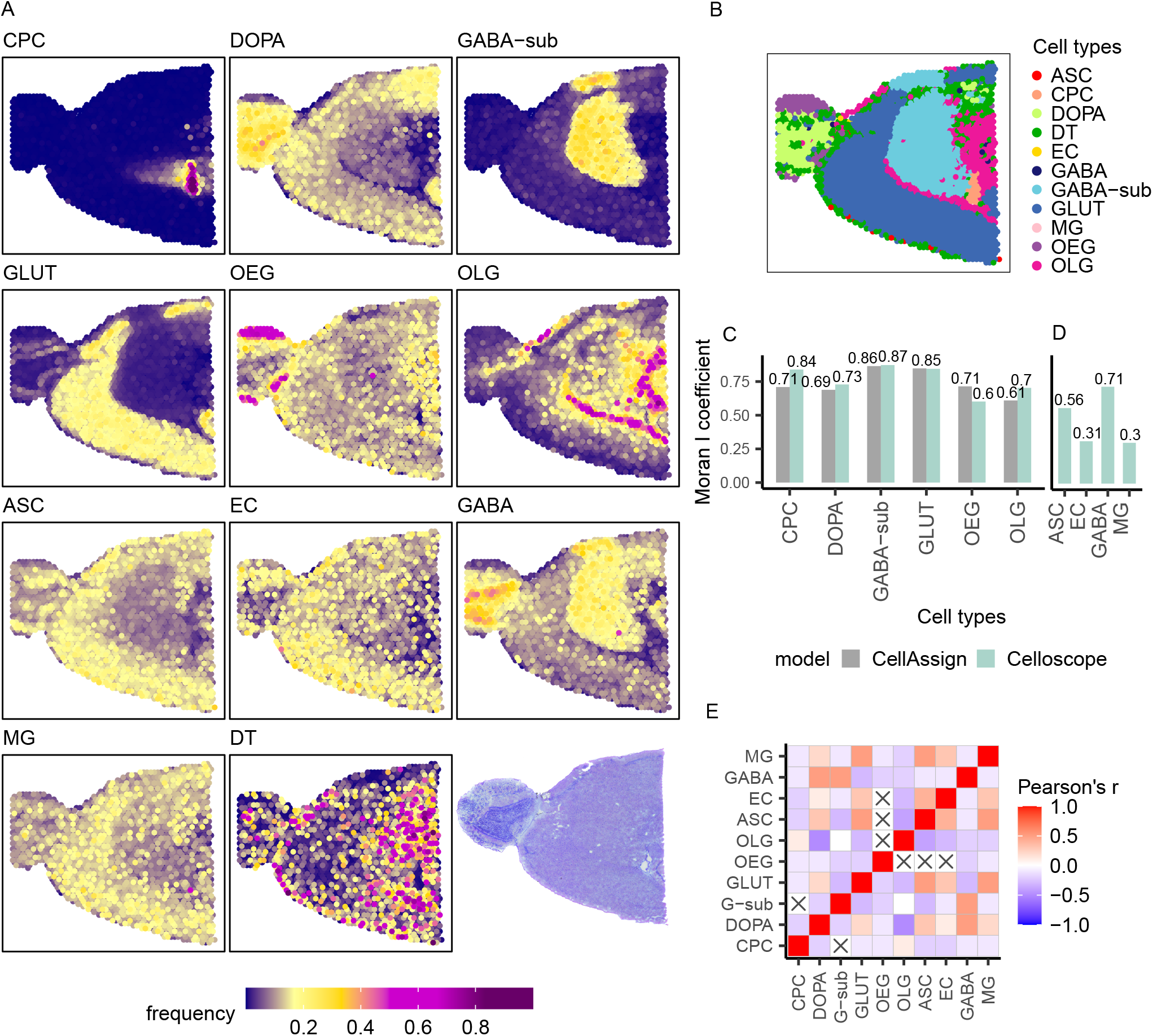
Results obtained for the anterior part of the mouse brain (sagittal section). OLG – Oligodendrocytes, OEG – Olfactory ensheathing glia, ASC – Astrocytes, GABA – GABAergic neurons, GABA-sub – GABAergic neurons subtype, GLUT – Glutamatergic neurons, DOPA – Dopaminergic neurons, CPC – Choroid plexus epithelial cells, EC – Endothelial cells, MG – Microglia, DT – dummy type. **A** Heatmaps represent spatial composition for selected cell types. Dark violet indicates the absence of the cell type in question, yellow signalises moderate occurrence and magenta domination of a given type. The last subfigure presents the corresponding H&E image. **B** Results of CellAssign on the same dataset. **C** Moran *I* coefficient for cell types indicated both by CellAssign and Celloscope. **D** Moran *I* coefficient computed for cell types indicated only by Celloscope. **E** The correlation matrix heatmap represents the values of the Pearson correlation coefficient for all studied cell types, the positive values in red, negative in blue. 0 indicates that there is no relationship between studied variables. “X” denotes an insignificant correlation (p-values of the test with the test statistics based on Pearson’s product moment correlation coefficient *p* ≤ 0.05).

In contrast to simulated data, for this dataset there is no ground truth specifying the exact, underlying compositions of cell types in each spot. We can, however, expect that some of the known cell types will dominate in specific brain regions and that some other non-specific cell types will be prevalent across the entire brain tissue. Thus, to evaluate the quality of cell type deconvolution by Celloscope, we first compared the obtained spatial cell type distribution to regions as specified by the mouse brain atlas (19). Second, we compared these findings to results of other studies localising cell types in mouse brain regions using different technologies, namely immunofluorescence detection (20), Nissl-sting for cells and genetic marker stains (21) or labeling with anti-TH antibody (22).

Neurons and non-neuron cells called glia are the most commonly occurring brain cells (23). Two major subclasses of neurons can be distinguished: GABAergic neurons establishing inhibitory synapses and glutamatergic neurons establishing excitatory synapses. We showed that Celloscope was able to spatially distinguish between the two of them: GABAergic neurons were found mainly in the olfactory bulb and olfactory cortex, while glutamatergic neurons were found mainly in the cerebral cortex (19). These results were similar to those obtained in (21), albeit using a different technology.

Glia do not produce electrical impulses, but rather provide support and protection for neurons. Celloscope identified that one of the main cell types of glial cells, astrocytes, which surround and support neuron functioning, are localized throughout the entire examined sample, similarly as found in (20). Microglia, the immune cells of the central nervous system, are constantly testing the environment for signals of malfunctioning and acting in the event of trouble. Their function justifies their omnipresence in limited quantities throughout the examined sample, as correctly identified by Celloscope.

Dopaminergic neurons synthesize the neurotransmitter dopamine. Similarly as in (22), these cells were found by Celloscope in the olfactory bulb. What is more, as expected, choroid plexus epithelial cells were localized in choroid plexus and olfactory ensheathing glia cells were found in the olfactory bulb.

#### Comparison to CellAssign

To show the benefits of accounting for the presence of cell type mixtures in spots as opposed to assuming they contain cells of single types, we compared the Celloscope’s outcomes to results obtained with CellAssign (17). Since CellAssign was originally developed to assign types to single cells based on scRNA-seq data, applying this approach to ST data is equivalent to considering each spot as homogeneous with respect to cell types, as if each spot was a single cell. Similarly to Celloscope, CellAssign correctly delineates mouse brain regions, assigning spots to dominating cell types for each region (Fig. 3B). In contrast to Celloscope, however, CellAssign per construction cannot identify cell types that are present in the examined tissue in lower prevalence, such as astrocytes, endothelial cells and microglia. For instance, while Celloscope indicates that astrocytes tend to occur in low amounts, mostly in the cerebral cortex, CellAssign omitted this cell type almost entirely, indicating only 11 spots out of 2696 to be dominated by astrocytes. Note that CellAssign was able to identify only a certain subtype of GABAergic neurons, while the more general type GABAergic neurons was disregarded and assigned to only seven spots. Lastly, microglia were identified in only two spots and endothelial cells in two. On this account, we distinguished two groups of identified types: indicated both by Celloscope and CellAssign (Fig. 3D) and cell types that were identified only by Celloscope (Fig. 3E). In summary, the results obtained by Celloscope and CellAssign were in agreement, however the Celloscope’s inherent feature of accounting for cell type mixtures enables it to provide a more comprehensive and more insightful description of the cell type composition of the tissue in hand.

#### Spatial autocorrelation of cell types

Given the naturally occurring tissue organization and structure, it is expected that neighboring spots will display spatial similarity and cells of the same type will co-localize. Note that Celloscope treats all spots as independent, regardless their position and potential proximity. As a consequence, spatial correlation across spots is not enforced in the model and can be used to validate the model’s performance. To this end, we calculate the Moran *I* coefficient (24, 25) to quantify the level of spatial autocorrelation of inferred cell type proportions (Fig. 3C, 3D). The Moran’s *I* coefficient takes values from –1 to 1, where –1 indicates perfect dispersion, 0 perfect randomness (no autocorrelation) and 1 perfect clustering of homogeneous values. Therefore, high values of the Moran *I* coefficient indicate that the inferred cell types cluster in space. We observe very high spatial autocorrelation for the majority of cell types and moderate for microglia and endothelial cells, however in all cases the obtained spatial autocorrelation is non-negligible. Note that the level of spatial autocorrelation for Celloscope is similar to autocorrelation level obtained by CellAssign, despite the fact of solving a more demanding and cumbersome task of cell type deconvolution as opposed to assigning the dominant cell type to a spot. Importantly, those cell types that were found in the tissue only by Celloscope and not by CellAssign also show spatial autocorrelation.

#### Spatial co-occurrence and mutual exclusivity between cell types

The cell type composition of spots resolved by Celloscope allows investigating spatial co-occurrence and exclusivity of cell types (Fig. 3E). We find that GABAergic neurons tend to spatially co-occur with dopaminergic neurons and GABAergic neurons subtype, microglia with astrocytes and glutamatergic neurons with astrocytes. On the other hand, Celloscope results suggest that oligodendrocytes avoid co-locallizing with dopaminergic neurons.

### Celloscope elucidated the source of inflammation in a human prostate tissue

Next, we applied Celloscope to analyze human prostate data (16). Again, investigation using the Gelman-Rubin method indicated that the sampler converged (Additional File 1: Figure S1 B). The analyzed dataset contained twelve sections from different regions of a resected prostate, which were profiled using ST. Several of these sections contained cancerous tissue. We selected one of the sections (3.1; Fig. 4B), where an immense infiltration of immune cells was visible in its respective H&E image. We applied Celloscope to investigate whether this infiltration could be associated with ongoing tumorigenesis in this area, as was observed by (26, 27), or whether it was due to some other inflammatory process. Notably, this information could not be derived from the H&E image alone, as the fine subtypes of the detectable cells were not distinguishable visually. For example, mononuclear cells with abundant, *foamy* cytoplasm indicating macrophages or cells with multilobed nucleus indicating neutrophils could be detected in H&E, but it was not possible to distinguish subtypes of lymphocytes (eg., T cells, B cells or NK cells). Similarly, raw gene expression data, measured using ST in this area, was not directly indicative of the type of the visible inflammation. Since detailed dissection of infiltrating immune cells identity is not feasible with classical histopathological inspection, nor directly from ST measurements, computational tools, such as Celloscope, are required to fill this gap.

**Fig. 4.**
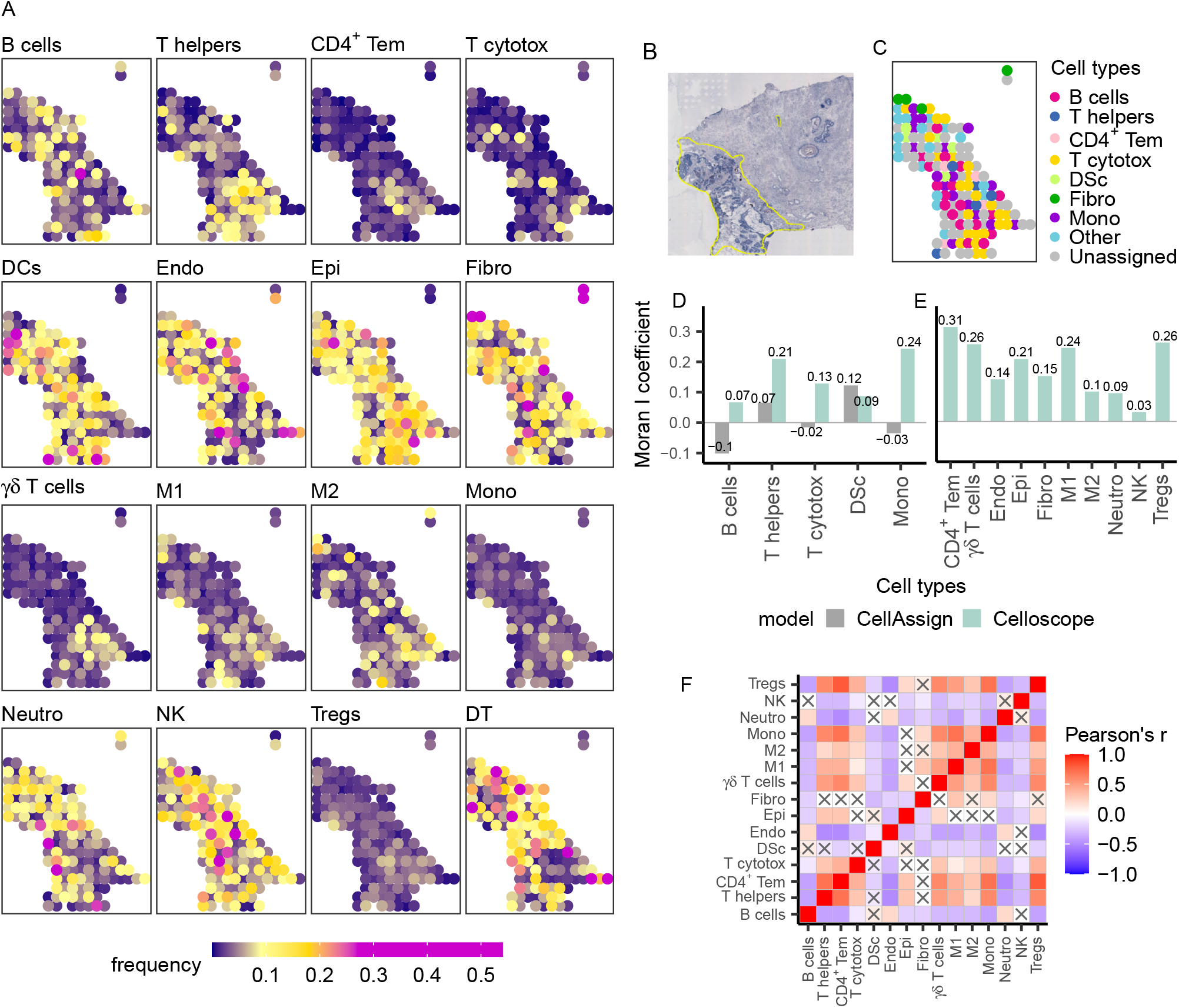
Results obtained for data from the human prostate. T helpers – CD4^+^ T cells (helper T cells), CD4^+^ Tem – CD4^+^ effector memory T cells, Cytotox – cytotoxic CD8^+^ T cells, DCs – dendritic cells, M1 – M1 macrophages, M2 – M2 macrophages, Mono – monocytes, Neutro – neutrophils, NK – Natural Killer cells, Tregs – natural CD4^+^ regulatory T cells, Endo – endothelial cells, Epi – epithelial cells, Fibro – fibroblasts, DT – dummy type. **A** Heatmaps represent spatial composition of cell types across spots. Dark violet indicates the absence of the cell type in question, yellow signalises moderate occurrence and magenta – domination of a given type. **B** The inflamed region of interest annotated on the H&E image (yellow selection). **C** Results of CellAssign performance. Colors correspond to cell types. Other – cell types that were indicated by CellAssign in less than 4 spots, namely: endothelial cells, epithelial cells, fibroblasts, γδ T cells, M1 macrophages, M2 macrophages, neutrophils, natural killer cells and CD4^+^ regulatory T cells. Unassigned – spots for which CellAssign was not able to indicate their type. **D** Moran *I* coefficient computed for cell types indicated both by CellAssign and Celloscope. **E** Moran *I* coefficient computed for types indicated only by Celloscope. **F** The correlation matrix heatmap represents the values of the Pearson correlation coefficient for all studied cell types, the positive values in red, negative in blue. 0 indicates that there is no relationship between studied variables. “X” denotes an insignificant correlation (p-values of the test with the test statistics based on the Pearson’s product moment correlation coefficient *p* ≤ 0.05).

To resolve the immune cell composition in this region, we aimed at identifying the following immune cell types across spots: B cells, CD4^+^ T cells (helper T cells), CD4^+^ effector memory T cells, cytotoxic CD8^+^ T cells, dendritic cells, γδ T cells, M1 and M2 macrophages, neutrophils, monocytes, natural killer cells and natural CD4^+^ regulatory T cells (Tregs). However, we also took into consideration non-immune cells that are expected to be present in the prostate tissue: endothelial cells, epithelial cells and fibroblasts.

Celloscope identified that the immune cell type composition of spots (Fig. 4A) is characterised by a larger heterogeneity as compared to the mouse brain tissue. In general, indicating the dominant type within each spot is more difficult, since for every spot (with only a few exceptions) the dominant type takes much less than a half of a spot capacity - the average most dominant type has the proportion of 21%. Moreover, several cell types (cytotoxic CD8^+^ T cells, monocytes and natural CD4^+^ regulatory T cells) are present only in trace amounts.

Celloscope identified the expected mutual arrangement of cell types such as general co-existence of epithelial, endothelial and stromal cells (fibroblasts) which compose the prostate gland. Endothelial cells, as expected, were found in the regions of edema, as well as in areas dominated by blood vessels. Fibroblasts are enriched in the regions without edema. Infiltrating immune cells, as expected, were found to be enriched in the edema regions. The found various immune cell populations (e.g. CD4^+^ helper T cells, γδ T cells, CD8^+^ cytotoxic T cells, macrophages, NK cells and neutrophils) were in agreement with previously observed highly heterogeneous composition of the immune infiltrate in the inflamed prostate (28, 29). Among immune cells identified by Celloscope, neutrophils, NK cells, and dendritic cells predominated and their dispersed distribution was in agreement with the morphological features characteristic for acute inflammation.

The composition of cell types identified using Celloscope allowed us to conclude that in this particular region of the resected prostate, the inflammation was not caused by tumorigenesis, but rather by an infection. As the examined region was the periurethral part of the prostate, the acute inflammation was most likely caused by a bacterial infection originating from the urethra. Identification by Celloscope of both neutrophils and NK cells in this area strongly suggests mixed infection with extra- and intracellular bacteria.

#### Comparison to CellAssign

We compared Celloscope’s outcomes to results obtained using CellAssign (Fig. 4C). CellAssign ignored numerous cell types (epithelial and endothelial cells, fibroblasts, neutrophils, M1 and M2 macrophages, γδ T cells, CD4^+^ effector memory T cells, and natural CD4^+^ regulatory T cells) and assigned each to only four or fewer spots. For those cell types that were identified both by Celloscope and CellAssign, their inferred localizations differ between the two approaches. For example, Celloscope identified that B cells cluster in small fractions of spots in the upper part of the inflamed region, whereas the helper T cells are clustered in the lower part. In contrast, CellAssign found these cell types as scattered across the entire region. Moreover, for those cell types that were ignored by CellAssign, such as γδ T cells (Fig. 4A), Celloscope again indicated a clustering of spots containing those cell types.

#### Spatial autocorrelation of cell types

To systematically quantify the extent to which cell types found using Celloscope and CellAssign cluster in space, we applied the Moran *I* coefficient, assessing spatial autocorrelation (Fig. 4D, E). Since we expected spatial similarities across spots in proximity and neither Celloscope nor CellAssign assume that this is the case and treat the spots as independent, spatial autocorrelation provides the means for an independent validation for models’ results. We observed moderate spatial autocorrelation for the majority of cell types, which was consistently higher for Celloscope than for CellAssign for cell types identified by both methods (Fig. 4D). Interestingly, we observed higher autocorrelation for cell indicated by Celloscope and ignored by CellAssign (Fig. 4E).

#### Spatial co-occurrence and mutual exclusivity between cell types

Finally, we inspected patterns of spatial co-occurrence and mutual exclusivity of cell types found using Celloscope (Fig. 4F). We found that CD4^+^ helper T cells, CD4^+^ Tem, CD8^+^ T cells (cytotoxic), γδ T cells, regulatory T cells and monocytes tend to co-localise. The largest positive spatial correlation was observed between regulatory T cells and CD4^+^ Tem. Since monocytes can differentiate into macrophages, it is not surprising that we observed high correlation between monocytes and macrophages M1, as well as M2.

## Methods

### Analysed data

#### Mouse brain dataset

ST profiling of the anterior part of the mouse brain tissue (a sagittal section) was generated with the Visium technology (v1 chemistry) from 10x Genomics. Raw data can be downloaded from (18). The analyzed count matrix (30) is an output of the spaceranger pipeline and (31) was accessed via `SeuratData` package (32) (Additional File 1: Section S4). The dataset contains expression of 31053 genes in 2696 spots, given by read counts. 182 genes were selected and modeled as markers (see below). Across the spots, a minimum of 9, on average 106, and a maximum of 141 genes had non-zero expression. The minimal total expression per spot was 10, maximal was 13513, while average total expression was 703.5.

#### Human prostate dataset

The analyzed human prostate data was generated by Berglund et *al.* (16). The raw files generated with ST were preprocessed by the authors as described in (33). The data comprised twelve sections from different regions of a resected prostate affected by cancer. From the total of 625 spots on section 3.1, we selected and analysed a subset of 274 spots that were contained in an area annotated as inflamed by an expert pathologist based on the respective H&E image. 161 genes were selected and modeled as markers (see below). Across the spots, a minimum of 6, on average 93.3, and a maximum of 159 genes had non-zero expression. The minimal total expression per spot was 24, maximal was 5138, while average total expression was 719.5.

### Curated marker gene sets based on literature and expert knowledge

#### Marker genes for the mouse brain dataset

For the analysis of the mouse brain dataset, a selection of marker genes was defined based on a previous study by Ximerakis *et al.* (34). In this study, firstly, cell type-specific marker genes, which were previously described in the literature, were used to annotate clusters found in a scRNA-seq dataset. Next, expression profiles of all the considered cell types were prepared. All genes expressed in considered cell types were ranked as markers based on their *quality* as defined by Ximerakis *et al.* (34). We used this ranking for the selection of the initial set of marker genes for our study. We further selected such genes that were active in the analyzed mouse brain dataset, i.e., had at least an expression count of four in at least five spots. The final set of 182 marker genes (Additional File 2: Table S1) was fixed after additional selection of such genes from the ranking of Ximerakis *et al.* (34), whose expression appeared to be spatially co-localized with the top 3 makers for each cell type in this ranking.

#### Marker genes for the human prostate dataset

For the human prostate data, cell types expected to be present in an inflamed area of the prostate tissue in a proximity of a tumor were first identified by an expert immunologist. Next, the initial set of marker genes was curated based on literature (35–38) and expert knowledge on markers of those immune cell types. The final set of marker genes for each cell type (176 genes in total) was manually selected based on the spatial agreement of their expression across spots (Additional File 3: Table S2).

#### Annotation of areas of interest based on the H&E image for the human prostate sample

In order to assess the type of tissue present in each spot for the human prostate dataset, we annotated contiguous tissue regions using QuPath (39). The annotated tissue types were as follows: suspected cancer, cancer, immune cells: chronic inflammation, immune cells: acute inflammation and suspected acute inflammation. To obtain lists of spots per each annotated area, we overlapped spot coordinates with tissue annotations using a custom script in QuPath.

#### Cell counting

We estimated the number of cells in each spot using a dedicated, custom script extending upon the functionality of QuPath (39). First, areas occupied by the circular spots in the analyzed H&E images were identified by their coordinates and diameter. Then, cell nuclei placed within these circular spot areas were detected using QuPath’s inbuilt cell counting algorithm for H&E image analysis (function `WatershedCellDetection`). To assure the counted cells were correctly distinguished from the background noise in the images by the algorithm, we manually, carefully adjusted the algorithm’s parameters, so that the returned cell counts in randomly selected spots were in agreement with counting cells by eye. The cell counting procedure was performed for all spots on the mouse brain slide and for the pathologist-selected inflammation area for the human prostate dataset.

#### The Celloscope model for cell type deconvolution in ST data

Celloscope is a novel, probabilistic, hierarchical Bayesian model of gene expression in ST data that can be used for marker gene-driven estimation of the proportions of different cell types present at selected spots of the examined tissue (Fig. 1). Celloscope represents all variables of interest as random variables (see Fig. 1b for graphical representation of the model, Tab. 1 for the list of variables and Tab. 2 for the hyperparameters). Let *s* ∈ {*s*_1_, *s*_2_,…,*s_M_* } index spots. We assume *t_N_* cell types are present in the considered tissue, indexed by *t* ∈ {*t*_1_, *t*_2_,…,*t_N_* }.

**Table 1.** Observed, hidden and deterministic variables in Celloscope, their corresponding probability distributions and interpretation.

| Observed variables |  |  |
| --- | --- | --- |
| $C_{gs}$ | Negative Binomial | gene expression for gene $g$ in cell $c$ |
| $B_{gt}$ | - | indicator whether gene $g$ is a marker for cell type $t$ |
| Hidden variables |  |  |
| $\theta_{st}$ | Gamma | unnormalised abundance of cell type $t$ in spot $s$ |
| $\lambda_0$ | - | base gene expression level, shared across all genes and cell types |
| $\Lambda_{gt}$ | - | over-expression for a marker gene $g$ in its specific cell type $t$ |
| $N_s$ | Truncated Normal | number of cells in spot $s$ |
| $p_g$ | - | success probability parameter for gene $g$ |
| $Z_{st}$ | Bernoulli | indicator variable stating presence of cell type $t$ in spot $s$ |
| $\pi_{st}$ | Beta | prior for the probability of presence of cell type $t$ in a spot $s$ |
| Deterministic variables |  |  |
| $h_{st}$ | Dirichlet | fraction of cell type $t$ in a spot $s$ |
| $\mu_{gs}$ | - | average expression of gene $g$ in a spot $s$ |

**Table 2.** Description of hyperparameters of Celloscope’s variables.

| Hyperparameters |  |
| --- | --- |
| $\alpha$ | adjustment for the average number of cell types present in a spot |
| $a$ | rate parameter for cell types present in a spot |
| $b$ | scale parameter for cell types present in a spot |
| $a_0$ | rate parameter for cell types absent in a spot |
| $b_0$ | scale parameter for cell types present in a spot |
| $l_s$ | estimate for the total number of cells in a spot |
| $\sigma$ | prior strength for $l_s$ |

Each type is represented by the set of marker genes. Those sets are potentially overlapping, but ideally, to better differentiate between the cell types, they should be disjoint. Let *g* ∈{*g*_1_, *g*_2_,…*g_K_*} be a set of marker genes. A binary marker gene signature for each cell type is encoded in a matrix *B*, such that an entry *B_gt_* takes the value 1 if a gene *g* is a marker for a type *t* and 0 otherwise. The matrix *B* is considered prior knowledge and is modeled as an observed variable.

The key assumption behind Celloscope is that marker genes are over-expressed in their specific cell types. To account for the increased expression of the marker genes in their specific cell types as compared to their base expression, we introduce the following two hidden variables: (i) Λ_*gt*_ as an over-expression of gene *g* when it is a marker for type *t*, and (ii) *λ*_0_ as a base expression of any gene shared across all the genes (for cell types other than specific for *g*).

Our main goal is to estimate the proportions of given cell types across all the spots, represented by the hidden variable *H*, which is a matrix with *s_M_* rows and *t_N_* columns. The value of an element *h_st_* is the proportion of all cells in spot *s* that are of type *t*, with values from 0 to 1. One row of the matrix *H*, denoted as *H*_*s*:_ = [*h*_*st*1_,…, *h_st_N__*] represents the hidden composition of spot *s*. Obviously, entries of a given row sum up to 1.

For a given gene *g*, its expression *C_gs_* is measured as the count of reads from spot *s* that map to gene *g*. The gene expression matrix *C* is modeled as an observed variable. A row *C*_*g*:_ = [*C*_*gs*_1__,…, *C_gs_M__*] represents the expression profile for gene *g* across cell types and a column *C*_:*s*_ = [*C_gs_*,…, *C_gK_ s*] represents expression profile for a spot *s* across marker genes. We assume that the expected value of the random variable *C_gs_* depends on the hidden composition *H*_*s*:_ of the cell types in spot *s*. In the following, we shall use the following three well known remarks:

#### Remark 1

*X* ~ *NB*(*r,p*) is equivalent to 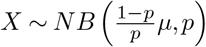, where 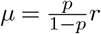.

#### Remark 2

Let *X,Y* be independent random variables satisfying *X* ~ *NB*(*r*_2_, *p*), *Y* ~ *NB*(*r*_2_, *p*). Then:

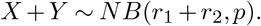

#### Remark 3

Let *X*_1_, …, *X_k_* be mutually independent random variables, *X_i_* ~ Gamma(*α_i_,β*), *i* = 1,…,*k* and 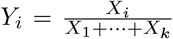. Then the joint distribution satisfies (*Y*_1_, *Y*_2_,…,*Y_k_*) ~ *Dirichlet*(*α*_1_, *α*_2_,…,*α_k_*).

Let us consider a single cell of an unknown type *t* present at the spot *s*. We denote expression of gene *g* in this cell as 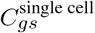. We use the Negative Binomial distribution to model gene expression. This distribution has two parameters: *r_gt_* - the rate parameter dependent on the gene and cell type in question, and *p_g_* - success probability dependent only on the gene in question. The variable *p_g_* enables to take into account the over-dispersion in gene expression data. We assume that:

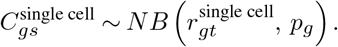

The average expression level of the gene *g* for that cell is equal to *λ*_0_ + *B_gt_*Λ_*gt*_. Thanks to Remark 1 we can express the rate parameter 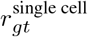 as the scaled mean of the considered distribution and we obtain:

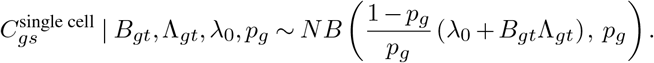

Obviously, a given spot *s* contains more than only one cell. Let us assume that at a given spot *s* there are *n_st_* cells of type *t*. Then we have

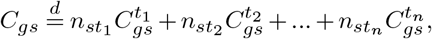

where 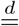 denotes equality in distribution.

Remark 2 gives us:

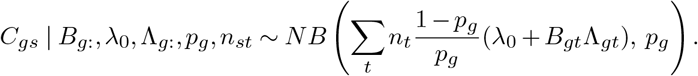

The total number of cells in spot *s* is represented by a hidden variable *N_s_*. We use the Truncated Normal distribution as the prior on *N_s_* with mean *l_s_* and variance *σ.* While *l_s_* is estimated based on H&E image analysis, the latter accounts for prior strength and the level of our belief in confidence in the results of number of cells estimation. Since *n_st_ = N_s_h_st_*, we have that:

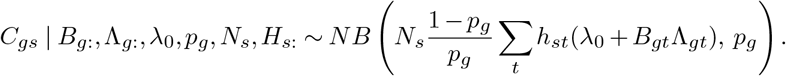

Denoting 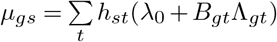, we obtain *t*

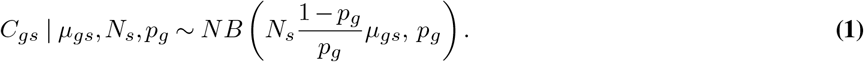

We use a simple feature allocation model to represent the presence of cell types in spots. We set

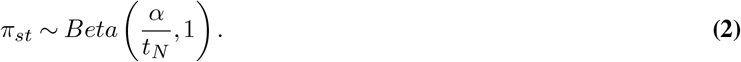

Let *Z_st_* indicate whether type *t* is present in spot *s* (takes value 1, if *t* is present in *s* and value 0 otherwise). We set:

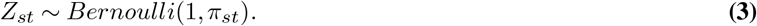

Let *θ_st_* denote the unnormalised abundance of type *t* in spot *s*. In the case when a given type *t* is present in spot *s* (*Z_st_* = 1) we expect *θ_st_* to take values more distant from 0 as compared to a situation when *Z* = 0. Therefore, we define a conditional distribution for *θ_st_* given the value of *Z_st_* in the following manner:

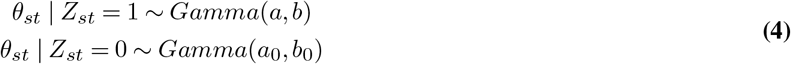

where *a, b, a*_0_, *b*_0_ are chosen so that values sampled from *Gamma*(*a, b*) are significantly larger than small values sampled from *Gamma*(*a*_0_,*b*_0_). Let us denote ⊖_*s*:_ = [*θ*_*st*1_, *θ*_*st*2_,…,*θ*_st_N__] as one row of matrix ⊖ describing unnormalised abundance of all cell types in spot *s*.

The proportion *h_st_* of cell type *t* in spot *s* is deterministically computed based on inferred ⊖_*s*:_. For a fixed spot *s*:

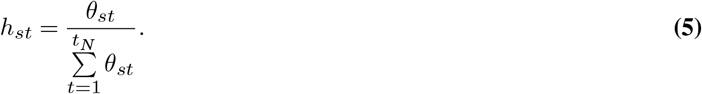

Remark 3 gives the conditional probability distribution of *H_s_*: given ⊖_*s*:_

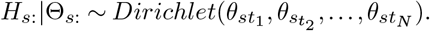

#### Celloscope’s parameters inference

For inferring the hidden variables we use the Metropolis-within-Gibbs sampler, a Markov chain Monte Carlo (MCMC) algorithm which is a combination of the Gibbs Sampler and the Metropolis-Hastings algorithm (Algorithm 1). Suppose the graphical model contains variables *x*_1_,…, *x_n_*. Let *MB*(*x_i_*) denote the Markov Blanket of *x_i_*, i.e., the set containing its parents, children, and co-parents. Then:

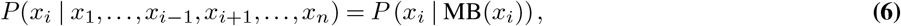

i.e., the conditional distribution of *x_i_* given the values of all other variables equals the conditional distribution given the values of the variables from its Markov blanket.

The iterative sampling procedure is as follows: firstly, starting values for 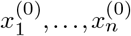 are randomly initialised, then, in a given iteration *j* (*j* = 1,2,…,*J*, where *J* denotes the number of iterations), we take every variable *x_i_*, *i* = 1,…,*n* one by one, in some arbitrary ordering and for each given variable *x_i_*, its value is sampled given the values of the variables from the Markov blanket 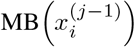 from the previous iteration. As a result each *x_i_* is updated iteratively, up until convergence. There are two options for updating the value of *x_i_* in the *j*-th iteration:

1. If 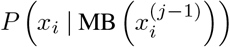can be expressed in a closed form, a new value 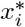 is sampled directly from 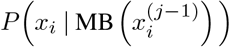 and 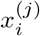 is set to 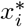.
2. In case we only known a function *f* proportional to *P* (*x_i_* | MB(*x_i_*))

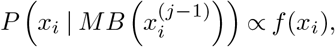

we perform a single Metropolis-Hastings accept-reject step (MH-single-step procedure in Algorithm 1). A candidate value 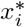 is sampled from a predefined proposal distribution 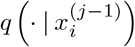, and then either accepted with probability given by

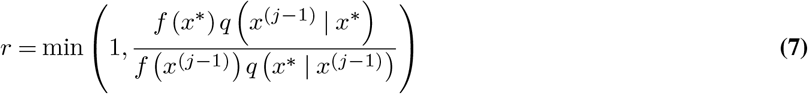

and 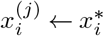, or the previous value is held: 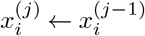. After updating *x_i_*, we immediately use the new value for sampling other variables.

**Algorithm 1.**
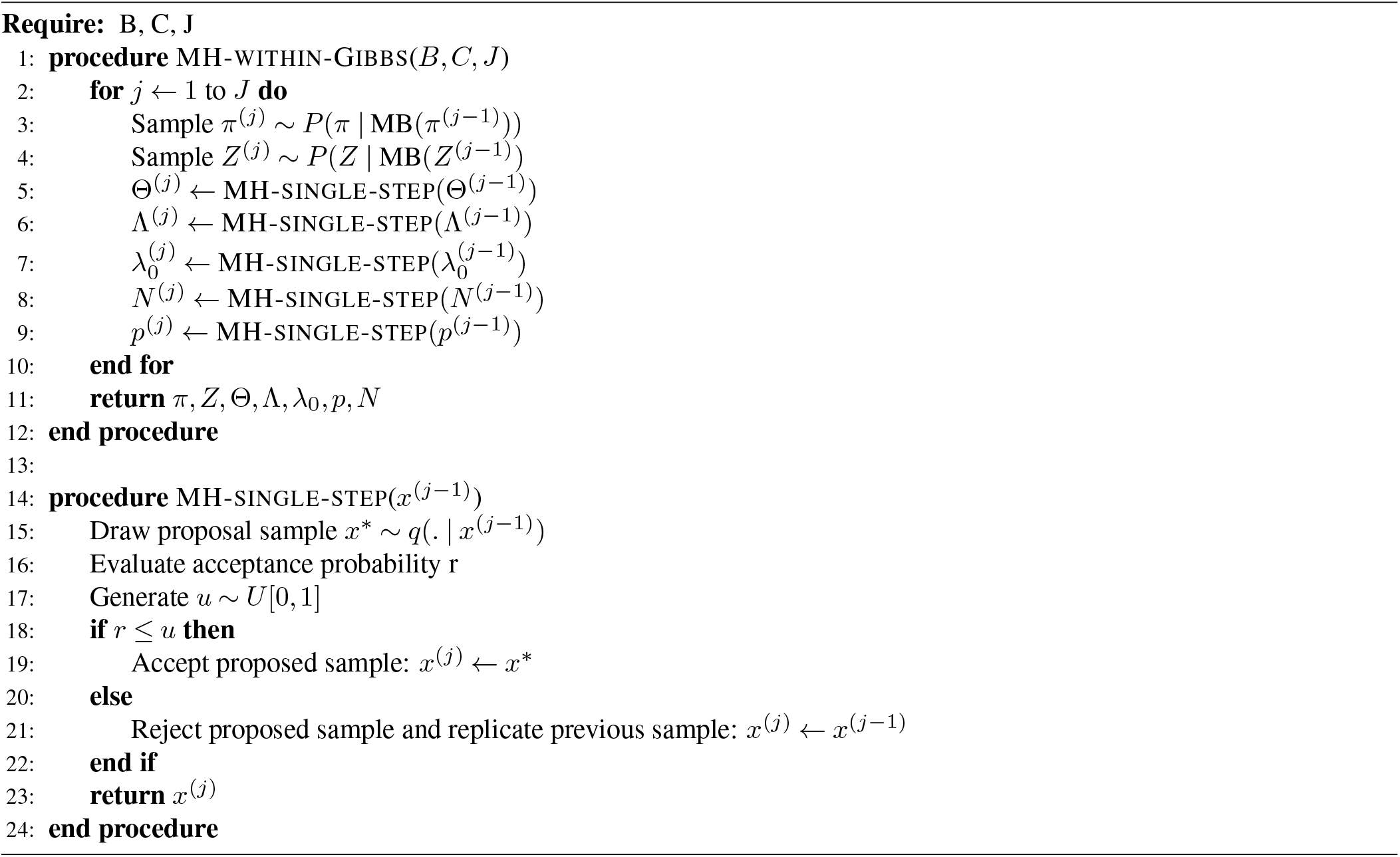
Metropolis-Hastings within Gibbs Sampler of Celloscope’s parameters

Below, we provide the model’s equations for target distributions, for which, in case of the first group of variables, namely *π* and *Z*, we know the explicit formulas. In case of the second group of variables: ⊖, Λ, *λ*_0_, *p*, *N* only functions *f* (·) proportional to the target distributions are known. These functions are used to compute the acceptance ratios (Equation 7), while performing single accept-reject Metropolis-Hastings steps.

##### Sampling π_st_

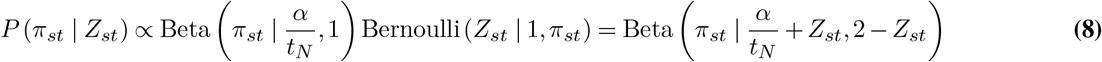

##### Sampling Z_st_

As *Z_st_* is a discrete, binary random variable, it suffices to consider its two possible values, 0 and 1.

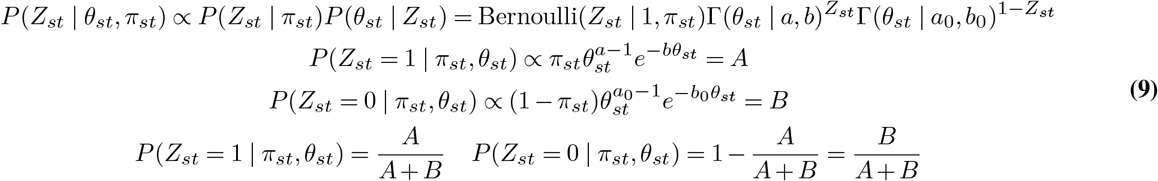

##### Target distribution for unnormalised cell type abundance θ_s_

We update each spot independently. For a selected spot *s*:

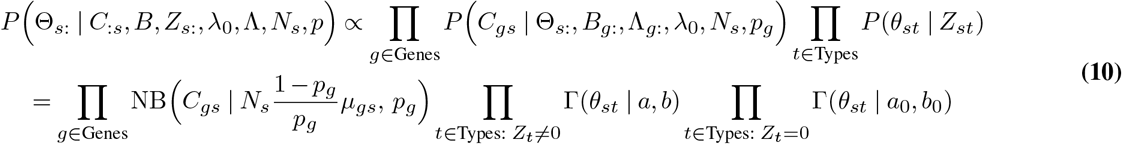

##### Target distribution for N_s_

For a selected spot *s*:

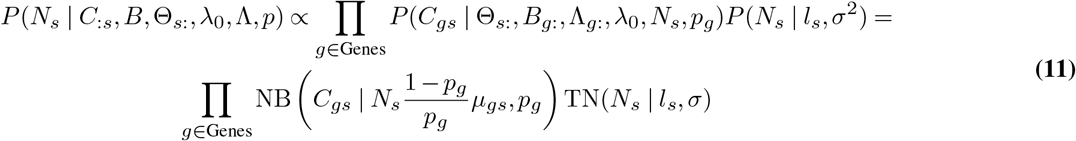

##### Target distribution for λ_0_

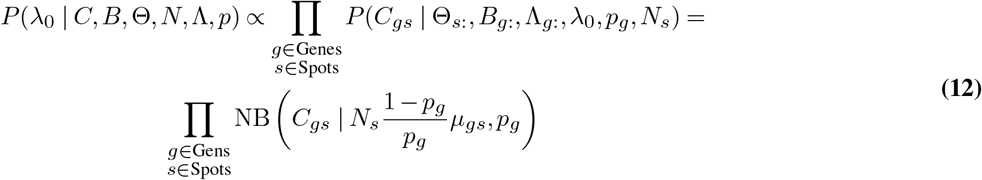

##### Target distribution for Λ_*gt*_

For a selected gene *g*:

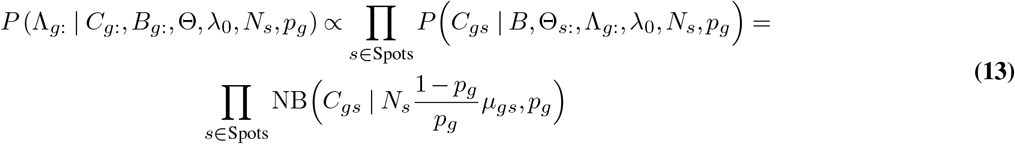

##### Target distribution for p_g_

For a selected gene *g*:

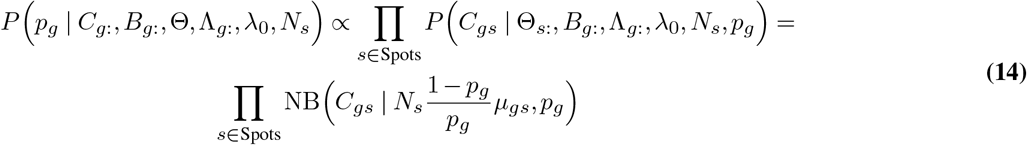

##### Proposal distribution

Let Φ(*x*) denote the cumulative distribution function of *N*(0,1), evaluated in point *x* and *q*(*x*|*y*) the proposal distribution, i.e., the conditional probability of proposing a new state *x* given the previous value was equal to *y*. We choose the truncated normal distribution *TN*(*μ,σ*) for the proposal distribution, since it allows for controlling the step size and it is appropriate for proposing values for non-negative variables. Note that for the truncated normal distribution we have that:

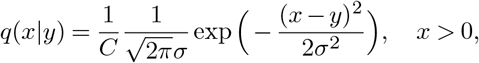

where 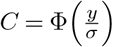 is a normalising constant. Bearing this in mind, we compute the Hastings ratio:

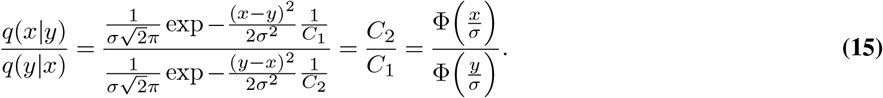

Equation (15) is used for all the model variables, except for *N_s_*, which unlike the rest, takes integer values. Therefore, we choose the ceiling of the truncated normal distribution as the proposal distribution. To compute the Hastings ratio in this case, we first need to find the density of a random variable denoting the ceiling of a truncated normal random variable.

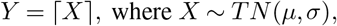

###### Remark 4

Let Φ_*TN*_ (*x*) denote the cumulative distribution function of *TN*(*μ, σ*^2^) in point *x*. Then:

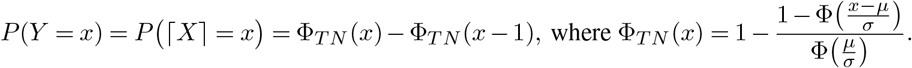

Proof:

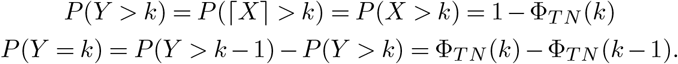

Now we can compute the Hastings ratio used for *N_s_*:

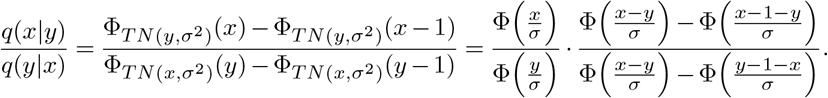

##### The dummy type

In order to account for cell types that potentially exist in the examined tissue but are unknown or not considered and thus are not represented in the cell type marker matrix *B* we introduce a so-called *dummy type*. We assume that the set of dummy type’s marker genes is separate from the set of all markers of the modeled cell types. Therefore, technically, to model the dummy type, we insert an additional column *t* for the dummy type and fill it with zeros, so that for all *g* ∈ Genes we have *B_gt_* = 0.

##### Adaptive step size

Let us recall that we employ the truncated normal distribution *TN*(·,*σ*) to propose a new value in the Metropolis-Hastings accept-reject step in Algorithm 1. The variance parameter *σ* corresponds to the *step size* of the sampler and affects the speed of convergence to the target distribution. To accelerate the convergence we modify the algorithm in such a way that we use a different step size for every sampled variable and adapt it as the sampling procedure progresses, aiming at achieving the optimal acceptance ratio of 23% (40). Specifically, for a selected model’s variable, we start with an arbitrary value for its step size, and once in every 5,000 iterations in case of mouse brain data and 10,000 in case of the human prostate data, we modify the step size according to the changing acceptance ratio. If the acceptance ratio is smaller than the optimal acceptance ratio, the step size is modified according to *σ* ← (1 – *e*)*σ*, otherwise *σ* ← (1 + *e*)*σ*, where *e* denotes a parameter controlling the strength of modification. Importantly, this procedure is restricted to the burn-in phase.

#### Synthetic data simulation scenarios

We considered four data simulating scenarios that differ with respect to the average number of distinct cell types present in each spot and the way the number of cells in each spot is accounted for in the model. In the first case, we either assume an increased number of cell types (dense scenario), or a decreased number of cell types (sparse scenario). While the dense scenario on average resulted in circa five cell types present in each spot, the sparse scenario resulted in circa two different cell types in each spot. When it comes to the second aspect, the way the number of cells in each spot is accounted for in the model, in the first option we use the number of cells in each spot in the inference as as the model’s input (**as known** values), i.e, *N_s_* variables become observed and their values are fixed to the true values. In the second option, we incorporate them as values for *l_s_* hyperparameters of the *N_s_* variables (as, so called, **priors**), i.e., the *N_s_* are hidden, inferred variables, and their prior mean is fixed to the true values.

#### Synthetic data simulation

We set *s_M_* = 800, *t_N_* = 8, *a* = 10, *b* = 1, *a*_0_ = 0.1, *b*_0_ = 1, λ_0_ = 0.2 and the number of marker genes *g_K_* = 149 (marker genes distributed across cell types as: 15,31,35,23,17,13,20). The value of the parameter *α* depends on the data simulating scenario, i.e. for the dense scenario we fix *α* = 2*t_N_* and for the sparse scenario *α* = 0.45*t_N_*. To sample values for gene expression level Λ, we firstly calculate average gene expression for cell types found in a scRNA-seq dataset on the mouse brain cortex (41). The obtained values were resampled to finally get values of Λ for all marker genes. The values for *p_g_* were sampled from *Unif* (0,1), *π* according to the distribution (2), *Z* according to distribution (3) and ⊖ according to (4). We computed *H* based on ⊖, from Equation (5).

After all values for hyperparameters have been established, for each of four considered scenarios, 15 replicates were sampled from the generative model Equation (1). As a result, we obtained 60 synthetic datasets.

#### Quality measure of the inference accuracy in synthetic data

For each of 60 synthetic datasets we calculate:

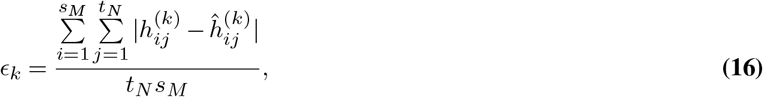

where *ϵ_k_* denotes the average error across spots and cell types for the *k*-th datasets, *k* = 1,…, 60, 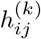 denotes the true value of a proportion of a type *j* in a spot *i* for the *k*-th dataset and 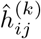 is the proportion of a type *j* in a spot *i* estimated by Celloscope for the *k*-th dataset, Stereoscope or RCTD.

#### Run setups for mouse brain and human prostate data

In the following, we describe the Celloscope’s run settings for the mouse brain dataset, and provide the corresponding settings used for the human prostate data in brackets. We ran three (ten) independent chains with the same hyperparameters (Tab. 3), but different (random) starting values with 100,000 (400,000) total iterations, including burn-in phase of the first 50,000 (300,000) iterations. The estimated total number of cells in each spot (*l_s_*) was incorporated as priors for the *N_s_* variable. For ⊖, Λ,λ_0_, *p_g_* we introduced adaptive step size (with the optimal acceptance ratio of 23%). Table 4 presents the starting values for the step sizes. Additionally, we set the thinning parameter to 10 (100), which resulted in keeping every 10th (100th) value and discarding the rest of them. Finally, estimates obtained via independent chains were averaged.

**Table 3.** Values of hyperparameters used for mouse brain data and human prostate data (in brackets).

| variable | hyperparameters |
| --- | --- |
| $\theta$ | $a = 8(10), b = 1, a_0 = 0.1, b_0 = 1$ |
| $\pi$ | $\alpha = 5(8)$ |
| $\sigma$ | 2(5) |

**Table 4.**
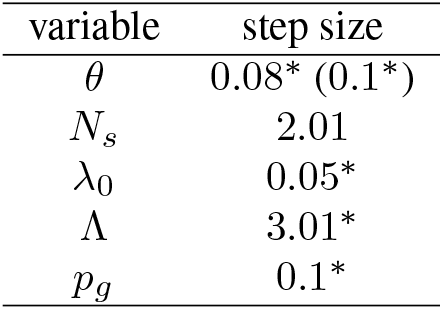
Step sizes’ values for proposal proposal distributions used to update variables with a single accept-reject Metropolis-Hastings step. * denotes adaptive step size.

| variable | step size |
| --- | --- |
| $\theta$ | 0.08* (0.1*) |
| $N_s$ | 2.01 |
| $\lambda_0$ | 0.05* |
| $\Lambda$ | 3.01* |
| $p_g$ | 0.1* |

## Discussion

In this contribution, we proposed a probabilistic Bayesian framework for comprehensive and accurate decomposition of cell type mixtures in ST spots based on marker genes. In this way, we enable spatial mapping of known cell types and indication of the presence of novel ones in tissues examined using the ST technology. Importantly, we are able to complete this task without a reference scRNA-seq data, as opposed to the efforts made hitherto. Thus, our method benefits from knowledge on marker gene sets that has been accumulated throughout many years, via different techniques and by independent researchers (15). Such incorporation of prior, domain knowledge was shown to limit the hypothesis spaces of models and guide them towards feasible solutions (42, 43).

The results of our extensive simulation study demonstrated the Celloscope’s accurate performance. Moreover, the outcomes of the conducted analyses of mouse brain data showed the Celloscope’s capability to unravel spots’s percentage composition with respect to cell types. Finally, the obtained insight into the source of inflammation in the human prostate dataset showed the Celloscope’s usefulness to investigate heterogeneous tissue’s functionality.The correctness and effectiveness of our approach is additionally demonstrated by the high spatial correlation for inferred cell type proportions measured with the use of the Moran *I* coefficient.

Not only has Celloscope numerous, significant advantages and sheer strengths, but also it substantially stands out from previously proposed methods that rely on integrating scRNA-seq with ST data. The fact that Celloscope is fully independent of scRNA-seq data intrinsically mitigates risks encountered while integrating data from the two disparate platforms. Further, we explicitly account for the total number of cells (regardless of their type) present at each ST spot by introducing a dedicated random variable to the model. This variable is particularly important as the number of cells may vary significantly across spots and extensively influence gene expression level measured for each spot. Therefore, accounting for the number of cells in each spot significantly boosts inference accuracy. Moreover, the Bayesian approach allows to incorporate additional information, such as the probability of a certain type being present at a certain spot, as priors.

Our results strongly advocate for the need to account for cell type mixtures in ST spots instead of assigning whole spots to single types. The advantage of accounting for mixtures was made evident by confronting our results with results obtained with CellAssign, a tool designed to assign types to single cells. On simulated data, CellAssign achieved much lower performance in terms of identifying the dominant type in spots. In application to prostate inflammation data, the Celloscope’s results are characterised by a significantly higher Moran *I* spatial correlation coefficient as compared to CellAssign.

There are however, certain limitations as far as applying Celloscope is concerned. The use of prior knowledge on marker genes naturally restricts the scope of considered cell types to a predetermined, closed list. To address this issue, we introduce the *dummy type* to account for the presence of other cell types than listed. As the Celloscope’s inference is being based solely on marker gene sets, our method requires human attention and expert knowledge, which may be demanding to acquire. Additionally, Celloscope’s ability to distinguish between cell types is limited to these cell types for which marker genes are determined, and, as a result, the rest of cell types may not be identified by Celloscope.

As the number of analysed high throughput sequencing datasets will grow, our current understanding of cell types and marker genes will expand even more. This will trigger the growth and strengthen the importance of marker-gene-driven methods such as Celloscope. In a broader context, let us notice that Celloscope can be perceived as a probabilistic framework for describing and modelling the signal conveyed by a mixture of different factors or entities (in our case cell types) and could be used in other deconvolution problems. In summary, Celloscope is a step forward in developing new probabilistic methods used to analyse spatial transcriptomics data and answer diverse biological questions.

## Availability of data and materials

Celloscope implementation and datasets supporting the conclusions of this article are available at https://github.com/szczurek-lab/Celloscope. The results were plotted using ggplot2 (44).

## Ethics approval and consent to participate

Not applicable.

## Authors’ contributions

A.G., E.S. and S.D.S. developed the probabilistic model. A.G. implemented the model and carried out the application of the model, supervised by E.S. A.G. and E.S. conceived the study. A.G. and E.S. wrote the paper. K.D. carried out the benchmarking of alternative methods. Ł.K., Ł.R and I.F. analysed the H&E images. A.G., D.N., Ł.K. and L.K. analysed and interpreted the model results. All authors provided critical feedback; helped shape the research and analysis; edited, reviewed and approved the manuscript.

## Additional Files

***Additional file 1 — Supplementary Information.*** Additional file 1 contains Supplementary Text and Supplementary Figures.

***Additional file 2 — Table S1.*** Marker genes for the mouse brain dataset.

***Additional file 3 — Table S2.*** Marker genes for the human prostate dataset.

