## Additional file 1: Supplementary Information for "Celloscope: a probabilistic model for marker-gene-driven cell type deconvolution in spatial transcriptomics data"

### Supplementary Text

#### S1 Running Stereoscope

We ran Stereoscope using the following command:

```
stereoscope run --sc_fit true_gene_expression.tsv p_g.tsv --st_cnt C_gs.tsv -ste 75000
    -stb 256 -lr 0.01 --gpu -o.
```

`true_gene_expression.tsv` contains true values for gene expression profiles for each gene in each cell type. `p_g.tsv` contains true values for the over-dispersion parameter $p_g$.

#### S2 Running RCTD

We ran RCTD using the following commands:

```
nUMI <- colSums(counts)
SpatialRNA_instance <- SpatialRNA(coords, counts, nUMI)
myRCTD <- create.RCTD(SpatialRNA_instance, reference, max_cores = 7, gene_cutoff = 0,
    fc_cutoff = 0, gene_cutoff_reg = 0, fc_cutoff_reg = 0)
myRCTD@cell_type_info$info[[1]] <- true_gene_expression
bulk = fitBulk(myRCTD)
sigma = choose_sigma_c(bulk)
my_results <- fitPixels(sigma, doublet_mode = 'full')
norm_weights = sweep(my_results@results$weights, 1,
    rowSums(my_results@results$weights), '/')
```

`counts` denotes ST counts matrix and `coord` denotes spots coordinates. Parameters: gene_curoff and fc_cutoff, hene_cutodd_reg and cutoff_reg were all set to 0, so that all marker genes were used for further cell type deconvolution. Since the mean expression was not calculated from scRNAseq data, but rather had to be given to the model directly, `cell_type_info$info[[1]]` was set to `true_gene_expression`, which consisted of true values for gene expression level for each gene in each cell type. This enables to perform the process of cell type deconvolution and cell type normalization without a reference scRNA-seq data set. However, to achieve this, the implementation had to be slightly adjusted (source codes available at `https://github.com/szczurek-lab/Celloscope`.

### S3 Running CellAssign

We ran CellAssign using the following commands:

```
calculated_sum_factors = calculateSumFactors(C_gs)
fit <- cellassign(exprs_obj = C_gs, marker_gene_info = B,
    s = calculated_sum_factors, learning_rate = 10^(-2), shrinkage = TRUE).
```

C_gs denotes gene expression matrix. B denotes a binary matrix with prior knowledge on marker genes. Learning rate was set to $10^{-2}$ for mouse brain data, $10^{-5}$ for human prostate data and $10^{-3}$ for simluated data.

### S4 Accessing mouse brain data

The count matrix was accessed via SeuratData [1] with the following commands:

```
InstallData("stxBrain")
brain <- LoadData("stxBrain", type = "anterior1")
```

More information: https://satijalab.org/seurat/v3.2/spatial_vignette.html.

### Supplementary Figures

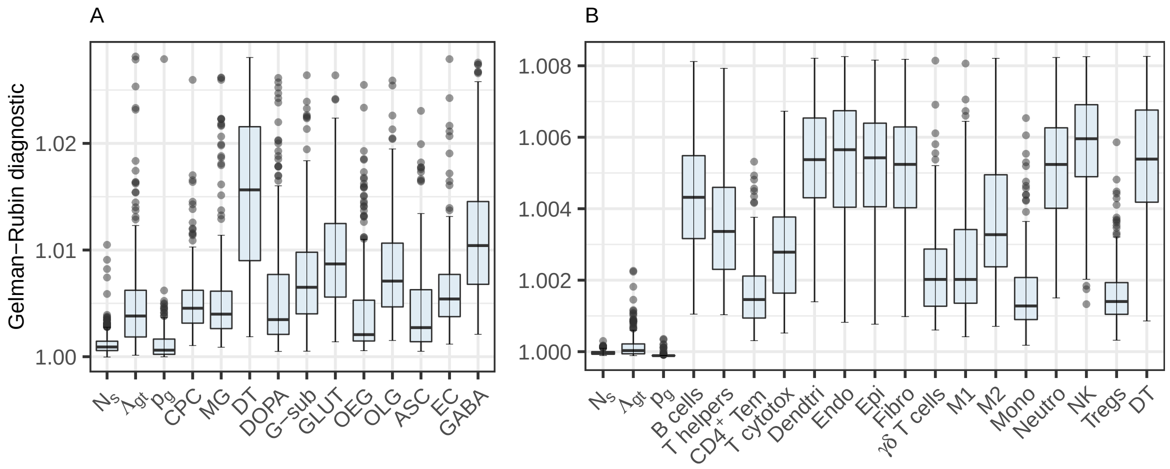


Figure S1: Box-plots represent values for Gelman-Rubin diagnostics test (R package stableGR [2]) with the division into model's variables for mouse brain (**A**) and human prostate data (**B**). In all cases the values are lower than the most commonly used convergence-indicating threshold: 1.1.
